# Gut microbiome communication with bone marrow regulates susceptibility to amebiasis

**DOI:** 10.1101/487652

**Authors:** Stacey L. Burgess, Jhansi L. Leslie, Md. Jashim Uddin, Noah Oakland, Carol Gilchrist, G. Brett Moreau, Koji Watanabe, Mahmoud Saleh, Morgan Simpson, Brandon A. Thompson, David T. Auble, Stephen D. Turner, Natasa Giallourou, Jonathan Swann, Zhen Pu, Jennie Z. Ma, Rashidul Haque, William A. Petri

## Abstract

The gut microbiome provides resistance to infection. However, the mechanisms for this are poorly understood. Colonization with the intestinal bacterium *Clostridium scindens* provided protection from the parasite *Entamoeba histolytica* via innate immunity. Introduction of *C. scindens* into the gut microbiota epigenetically altered and expanded bone marrow granulocyte-monocyte-progenitors (GMPs) and provided neutrophil-mediated protection against subsequent challenge with *E. histolytica*. Adoptive transfer of bone-marrow from *C. scindens* colonized-mice into naïve-mice protected against ameba infection and increased intestinal neutrophils. Because of the known ability of *C. scindens* to metabolize the bile salt cholate, we measured deoxycholate and discovered that it was increased in the sera of *C. scindens* colonized mice, as well as in children protected from amebiasis. Administration of deoxycholate alone (in the absence of *C. scindens*) increased the epigenetic mediator JMJD3 and GMPs and provided protection from amebiasis. In conclusion the microbiota was shown to communicate to the bone marrow via microbially-metabolized bile salts to train innate immune memory to provide antigen-nonspecific protection from subsequent infection. This represents a novel mechanism by which the microbiome protects from disease.

**One Sentence Summary:** Introduction of the human commensal bacteria *Clostridium scindens* into the intestinal microbiota epigenetically alters bone marrow and protects from future parasite infection.

## Main Text

Commensal intestinal bacteria can protect from infection (*1, 2*) in part by modulating bone marrow production of immune effector cells such as neutrophils and inflammatory macrophages (*3, 4*). In addition infection with one organism may persistently alter innate immune populations via microbial metabolites to provide protection from infection with unrelated pathogens in a process coined “*trained immunity*” (12, 13). Epigenetic changes in genes important in innate immunity have been implicated as a mechanism for trained immunity (*5–8*). These include changes in histone H3K27 and H3K4 methylation associated with promotor regions of innate inflammatory genes (*7*). As such, microbial metabolite alteration of H3K27 demethylase expression might contribute to the development of innate trained immunity (*7, 9, 10*). Host damage-associated molecular pattern molecules (DAMPs) that can be systemically induced by the microbiota have also been shown to be important in upregulating demethylase expression in both myeloid cell lines and mouse bone marrow (*11, 12*). Collectively, these data support a role of serum soluble mediators induced by the microbiota in communicating to the bone marrow to influence immunity to infection. We sought here to better understand the mechanism by which protective trained immunity by the microbiota might occur during a human intestinal infection.

We had previously shown that gut colonization with the *Clostridia*-related mouse commensal Segmented Filamentous Bacteria (SFB) protected from *Entamoeba histolytica* infection (*13*). SFB persistently expanded bone marrow granulocyte monocyte progenitors (GMPs) (*12*) which produce neutrophils which are known to protect from amebiasis (*14–16*). We hypothesized that components of the gut microbiota might alter bone marrow hematopoiesis to confer protection against an unrelated pathogen such as Entamoeba (*17, 18*). To explore this possibility we first tested for human commensals associated with protection from amebiasis (*19*). Principal coordinate analysis of beta-diversity indicated that the microbiome of children with *E. histolytica* diarrhea differed significantly (Figure 1 A) with a decrease in the relative abundance of the genus *Lachnoclostridium* (Figure 1 B). *Lachnoclostridium* have a number of members known to alter the metabolome including the bile acid pool of the intestine (*20*). We hypothesized that these bacteria may provide protection from ameba in part by altering the serum bile acid pool. To test this hypothesis we introduced the human commensal *Lachnoclostridium* related bacteria *Clostridium scindens* (*21, 22*) into the gut microbiome of susceptible CBA/J mice (*23*) and challenged them with the parasite *E. histolytica*.

**Figure 1.**
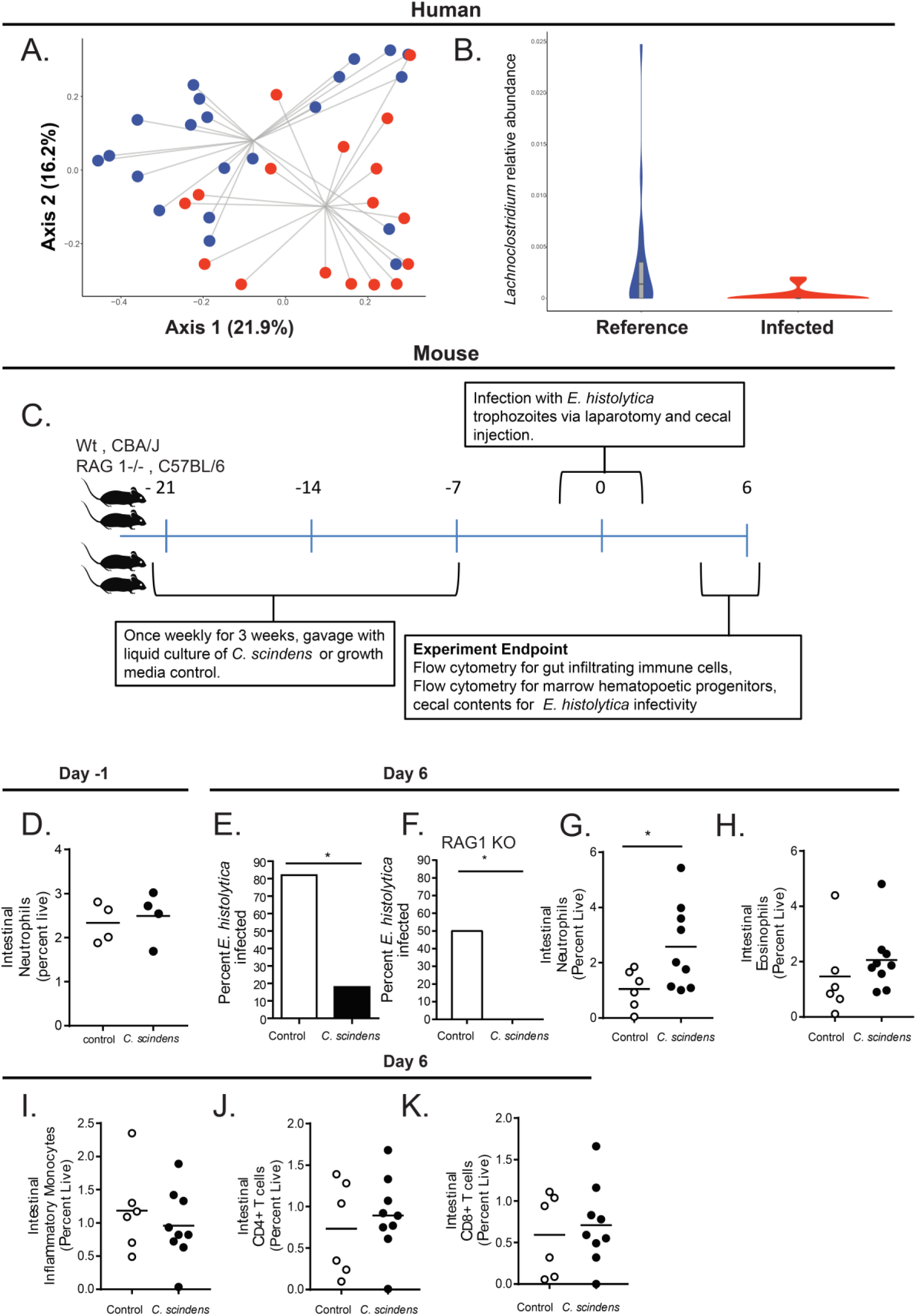
*Lachnoclostridium* are associated with protection from *Entamoeba histolytica* and introduction of *Clostridium scindens* to the gut microbiota provides innate protection from *Entamoeba histolytica* in a murine model. **(A)** Principal Coordinate Analysis (PCoA) of Bray-Curtis dissimilarities (beta-diversity) of fecal microbiota from surveillance reference stool or *E. histolytica* infected children. The groups are significantly different by PERMANOVA. **(B)** Relative abundance of the genus Lachnoclostridium from samples described in **A**. The groups are significantly different by Wilcoxon rank sum test with continuity correction. **(C)** CBA/J mice or C57BL/6 RAG1 KO mice were colonized with bile acid 7α-dehydroxylating bacteria *C. scindens* (ATCC® 35704) over three weeks prior to intracecal infection with *E. histolytica.* (D) Gut neutrophil infiltration was determined prior to ameba infection via flow cytometry. **(E, F)** Percent of mice infected with Entamoeba at day six following infection was determined via cecal culture in trophozoite culture media. **(G-K)** Gut immune cell infiltration was determined via flow cytometry. * p<0.05, PERMANOVA (ordination), Student’s t-test, Mann–Whitney U test, bars and error bars are mean and SEM. p= 0.006365, Wilcoxon rank sum test. N = 4-9 mice per group N= 20 children per condition.

*C. scindens* was significantly increased in the microbiota after gavage as measured by relative expression of the baiCD oxidoreductase, and gut community structure was also altered (Figure S4, A, B). Introduction of *C. scindens* to the gut microbiome provided protection from *E. histolytica* (Figure 1 C, E, F, Figure S3, S5) and this protection was associated with increased intestinal neutrophil infiltration (Figure 1 G). This increase in gut neutrophils only occurred with Entamoeba infection (Figure 1 D). There was no significant difference in intestinal CD4+ and CD8+ T cells, eosinophils or inflammatory monocytes (Figure 1 H-K) in *C. scindens* colonized mice. Gavage with a non-*Clostridia* human mucosal anerobic bacteria did not induce protection from Entamoeba (Figure S5).

Myeloid cell expansion such as observed here may be influenced by cytokine production by CD8+ T cells (*24*) or intestinal T regulatory cell accumulation (*25*). To test for a contribution of the acquired immune system to *C. scindens-*mediated protection we utilized RAG-1^-/-^ mice which lack B and T cells. RAG-1^-/-^ mice were also protected from *E. histolytica* when colonized with *C. scindens* (Figure 1 F) indicating that protection did not require the acquired immune system.

The increase in gut neutrophils in response to *Entamoeba* infection in *C. scindens* colonized mice suggested that *C. scindens* may have altered innate bone marrow populations that give rise to neutrophils. Therefore we examined hematopoietic progenitors in *C. scindens* colonized specific pathogen free mice (SPF) (Figure 2 A, B), SPF RAG-1^-/-^ mice (Figure 2C) and *C. scindens* gnotobiotic mice and germ free controls (Figure 2 D). Intestinal colonization with *C. scindens* increased bone marrow granulocyte progenitor cells (GMPs, CFU-GM) (Figure 2 A, B). Expansion of GMPs mediated by *C. scindens* occurred in the absence of T cells (Figure 1 C) and colonization with *C. scindens* alone was sufficient to increase marrow GMPs (Figure 1 D). This suggested that innate immune cells primarily underlie the observed *C. scindens* mediated changes in hematopoiesis and protection from *Entamoeba*. Furthermore the work suggested there may be a persistent epigenetic change in the GMPs that could support increased neutrophil production with *Entamoeba* challenge and facilitate bone marrow mediated protection from the parasite. To explore this possibility we examined transcriptional and epigenetic changes in sorted marrow GMPs from *C. scindens* colonized mice (Figure S2A).

**Figure 2.**
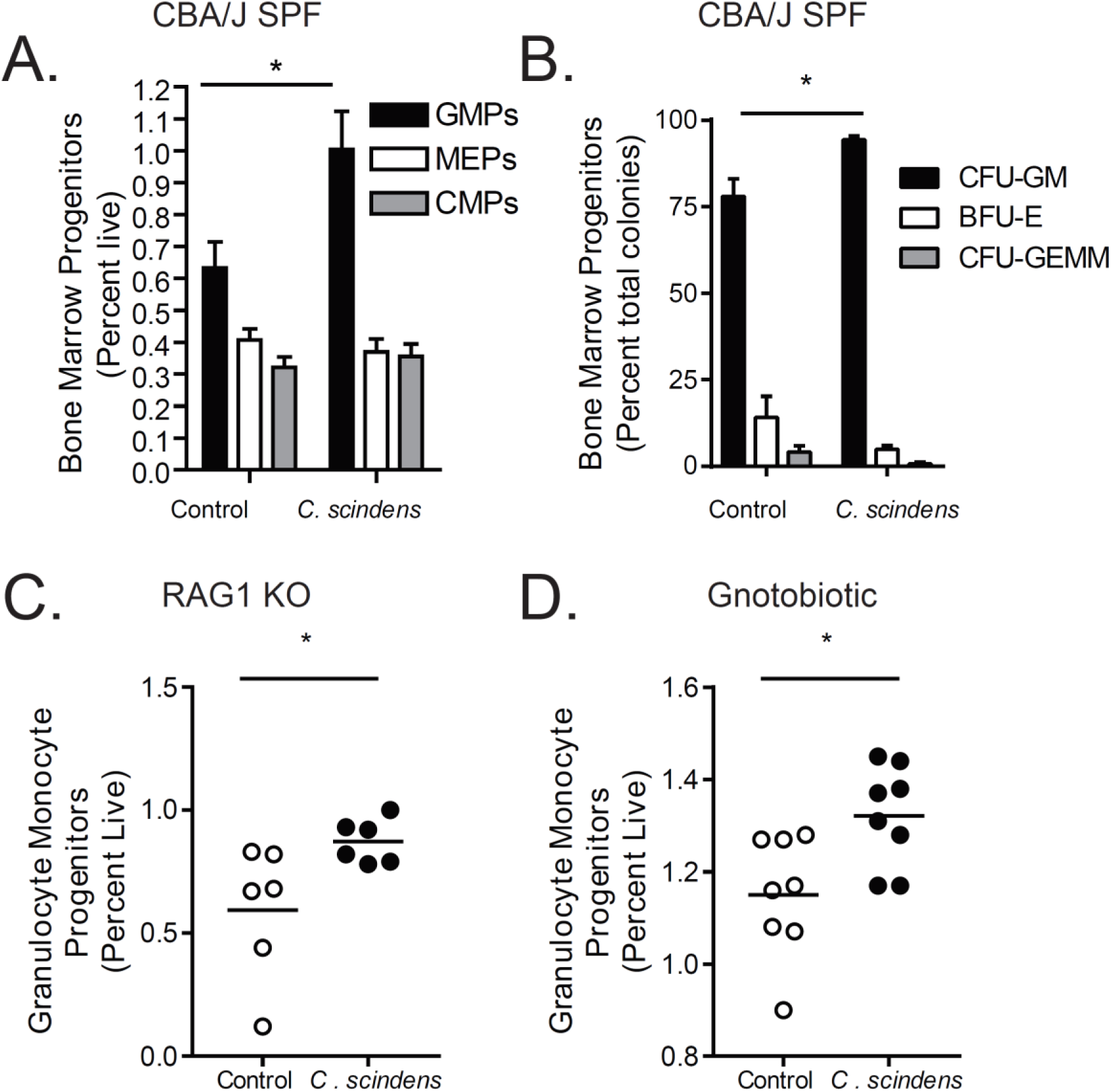
Intestinal colonization with *C. scindens* expands bone marrow granulocyte monocyte progenitors. CBA/J mice or C57BL/6 RAG1 KO mice were colonized with bile acid 7α-dehydroxylating bacteria *C. scindens* (ATCC® 35704) over three weeks prior to intracecal infection with *E. histolytica.* **(A, C, D)** Flow cytometry and **(B)** colony forming assays were utilized to determine composition of marrow hematopoietic precursors in *C. scindens* colonized CBA/J or RAG1 KO mice. *= p<0.05, Student’s t-test, bars and error bars are mean and SEM. N =6-8 mice per group.

RNA sequencing and gene enrichment analysis suggested that genes associated with covalent modification of the histone H3 tail, such as the demethylase JMJD3, are enriched and upregulated in mice exposed to *C. scindens* (Figure S2 A, B, C). This includes the enrichment of genes associated with CCAAT/enhancer-binding proteins, known to be important for GMP and neutrophil differentiation and expansion (*26*) (Figure S2 B). QPCR of sorted marrow GMPs demonstrated that significant changes in expression occurred in JMJD3 and CEBPA (Figure S2 D-G, I) Therefore we examined H3K4me3 and H3K27me occupancy in the promoter regions of two genes from this analysis known to be important in granulopoiesis, CEBPA(*27*) and CEBPB, in sorted GMPs. The repressive mark H3K27me3 was decreased in the promoter of CEBPA in *C. scindens* colonized mice (Figure 3A) while the activating mark H3K4me3 (Figure 3B) was increased in the promoter of CEBPB in *C. scindens* colonized mice. This indicated that bone marrow epigenetic alteration occured with gut colonization of *C. scindens* and suggested that persistent bone marrow changes might underlie the increased gut immunity to Entamoeba in colonized mice. To explore this possibility we utilized adoptive marrow transplants.

**Figure 3.**
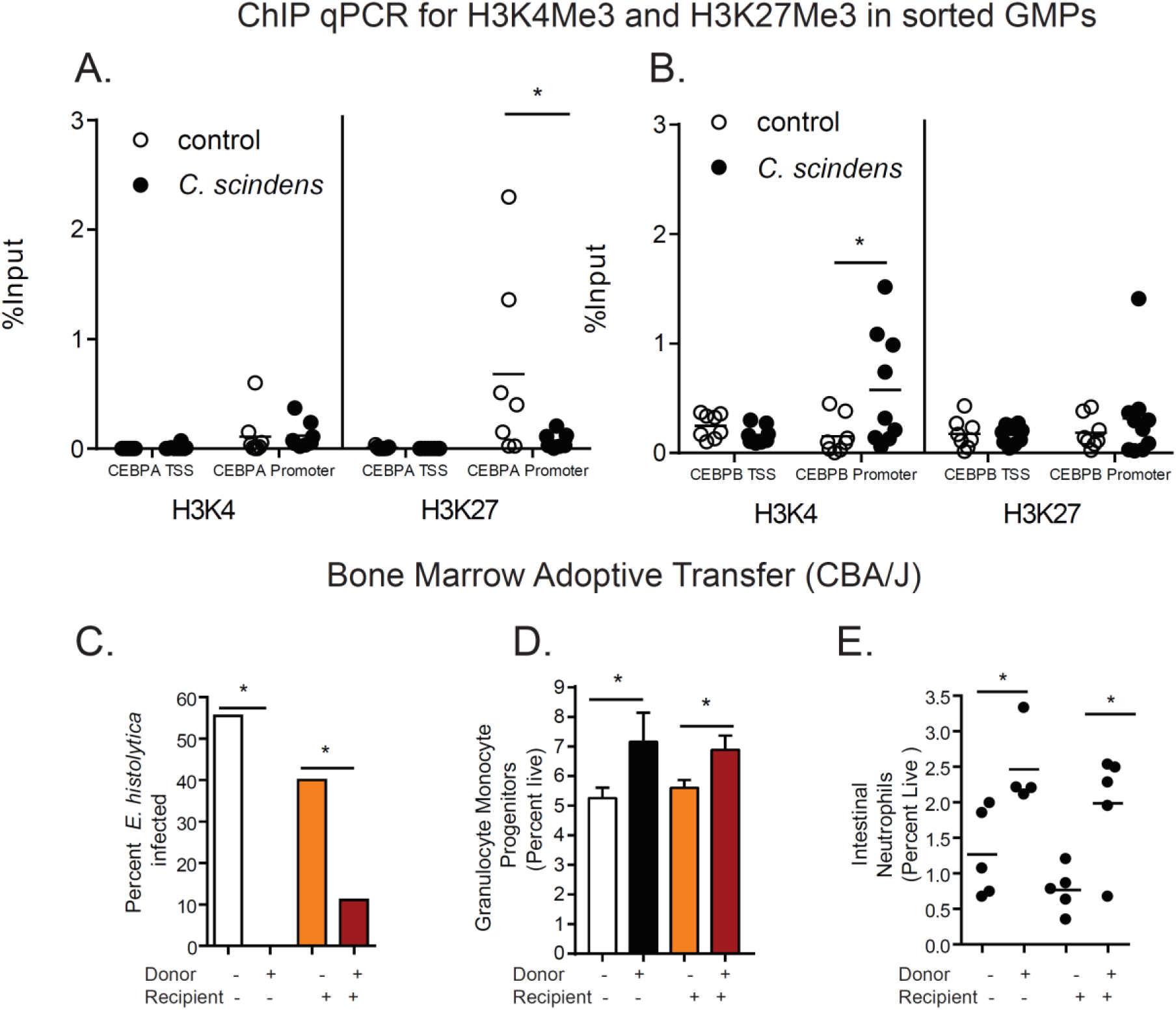
*Clostridium scindens* colonization epigenetically alters granuolocyte monocyte progenitors and bone marrow from *C. scindens* colonized donors is sufficient to provide protection from Entamoeba in *C. scindens* naïve mice. **(A, B)** ChIP for H3K4me3 and H3K27me3 was preformed followed by qPCR for the transcription start site (TSS) and promoter for CEBPA and CEBPB on sorted GMPs from mice colonized with *C. scindens* or control mice. **(C-E).** CBA/J mice colonized with *C. scindens* (+) or not (-) were lethally irradiated and given whole marrow from *C. scindens* (+) or *C. scindens* (-) donors then allowed to recover for 7 weeks prior to Entamoeba challenge. **(C)** Percent infectivity with ameba**, (D)** change in marrow GMPs, and **(E)** gut neutrophil infiltration were determined at 8 weeks post BMT. *= p<0.05, Student’s t-test, Mann–Whitney U test, One Way ANOVA with Tukey post-test, bars and error bars are mean and SEM. N = 6-8 mice per group.

Adoptive transfer of bone marrow from *C. scindens* colonized mice into mice not previously exposed to *C. scindens* was sufficient to provide protection from *E. histolytica* as well as recapitulate the observed increase in marrow GMPs and intestinal neutrophils. In contrast previous epithelial exposure to *C. scindens* was not sufficient to provide protection from ameba in irradiated mice (Figure 3 C-E). We concluded that long-lasting alterations in marrow hematopoietic cells caused by gut exposure to C*. scindens* were sufficient to confer protection via a more robust intestinal neutrophil response to later *E. histolytica* challenge.

We next explored how *C. scindens* could be epigenetically reprogramming GMPs in the bone marrow. *C. scindens* is capable of 7α-dehydroxylation of bile acids in the intestine (*4, 29*). As we expected, colonization of mice with *C. scindens* increased serum levels of the secondary bile acid deoxycholic acid (DCA, a product of of 7α-dehydroxylation of cholic acid) and increased bile acid deconjugation (Figure 4 A, Figure S1, Figure S2 H). Deoxycholate was also increased in children protected from *E. histolytica* (Figure 4 B). We replicated this finding that serum DCA independently predicted intestinal *E. histolytica* infection in a second childhood cohort from Bangladesh (Table ST1). We concluded that DCA in plasma was positively correlated with protection from Entamoeba in the mouse model of amebic colitis and in children.

**Figure 4.**
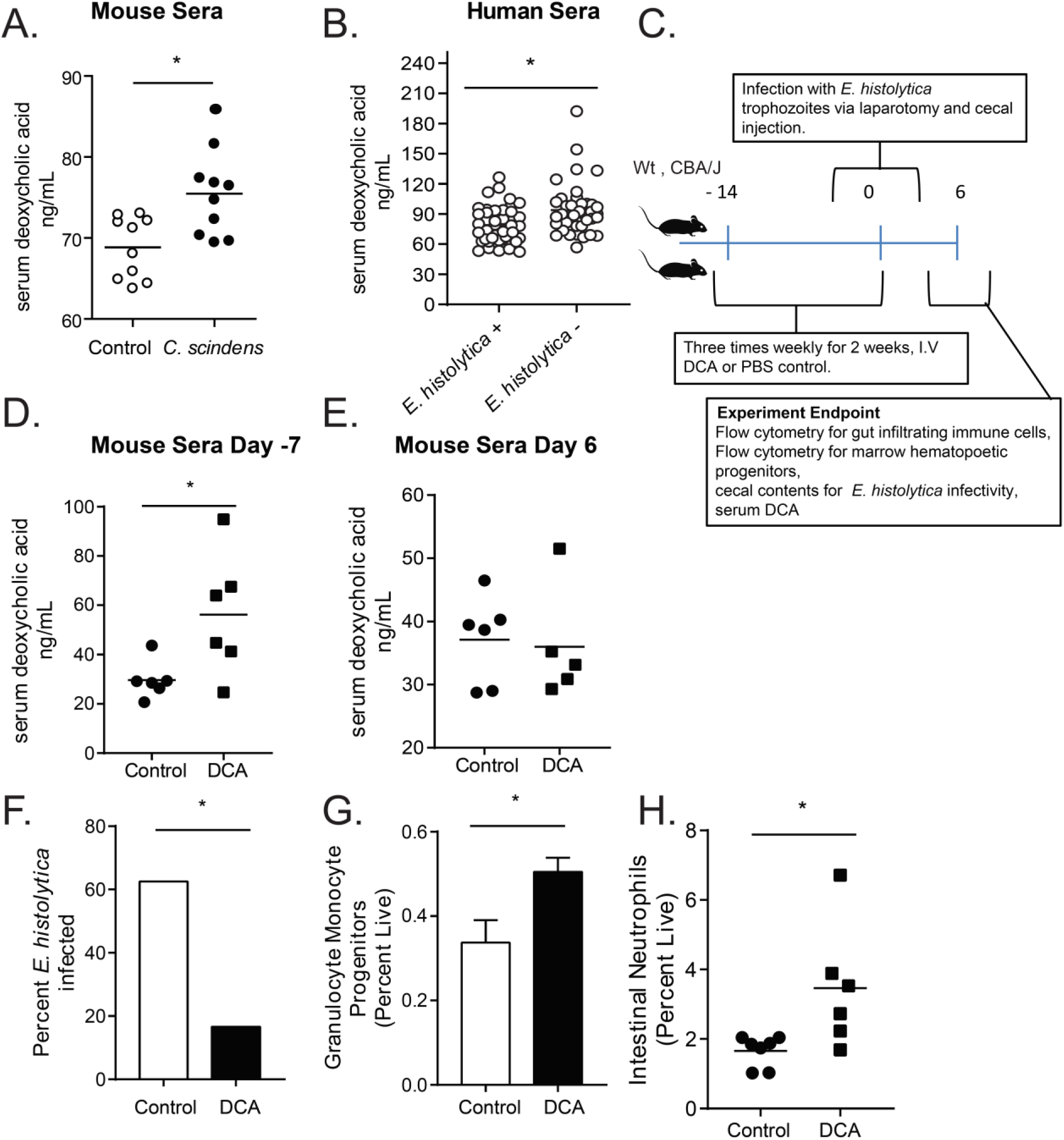
*C. scindens* colonization increases serum deoxycholic acid (DCA) and increased serum DCA is associated with protection from Entamoeba in both children and in a mouse model. **(A)** CBA/J mice were colonized with *C. scindens* and serum DCA was measured. **(B)** Serum DCA was measured via ELISA in 2 year old children in Bangladesh that were free of (-) or infected with (+) *E. histolytica* within 6 months of the blood draw**. (C).** Mice were administered DCA intravenously three times a week for two weeks and then challenged with *E. histolytica*. Serum DCA was measured during week 1 **(D)**, and at the end of the experiment **(E).** Percent infectivity **(F)**, change in marrow GMPs **(G)** and gut neutrophils **(H)** were measured at the end of the experiment. *= p<0.05, Student’s t-test, Mann–Whitney U test. N =6-8 mice per group. N= 40 children per condition.

To test if DCA was sufficient to mediate trained immunity against Entamoeba we administered the bile salt intravenously. Administration of deoxycholate prior to Entamoeba infection increased serum levels of deoxycholate to levels comparable to *C. scindens* colonization (Figure 4 C, D, and E) and provided protection from infection in the animal model (Figure 4 F). Protection from *Entamoeba* was associated with increased marrow GMPs and gut neutrophils (Figure 4 G, H). Additionally, experimental elevation of serum DCA increased expression of the epigenetic mediator JMJD3 in sorted marrow GMPs (Figure S2 G-I). We concluded that DCA was sufficient to replicate the changes in GMPs and protection from *Entamoeba* afforded by *C. scindens.* These studies however do not rule out the potential contribution of other bile acids and metabolites to gut to bone marrow communication.

Deoxycholate-mediated protection from *E. histolytica* was associated with increased marrow GMPs and intestinal neutrophils as seen with *C. scindens* (Figure 1 E, F, G). We were interested in pathways by which deoxycholate or *C. scindens* increased GMPs. Due to the epigenetic changes observed (Figure 3 A, B), persistent nature of immunity to *E. histolytica* following bone marrow transplant (Figure 3 C-E), and upregulation of JMJD3 in sorted marrow from *C. scindens* colonized or DCA treated mice (Figure S2), we examined the role of JMJD3 activity during *C. scindens* colonization on protection from *Entamoeba* infection. Treatment with an inhibitor of JMJD3 during *C. scindens* colonization abrogated bone marrow GMP expansion (Figure S3 A) as well as induction of intestinal neutrophils and protection from *E. histolytica* (Figure S3 E, F). This suggests that this epigenetic mediator and H3K27 demethylase activity may contribute to gut to marrow communication by *C. scindens*. Future studies will examine this possibility in more depth. JMJD3 is an H3K27me3 demethylase (*10, 30, 31*), however, we also observed changes in H3K4me3 in the promoter region of CEBPB. JMJD3 has recently been shown to impact H3K4me3 levels in human acute myeloid leukemia (AML) cells (*32*). However this may not fully explain the epigenetic changes in our model and other epigenetic mediators, including other non-methyl modifications such as H3K27Ac, might influence gut microbiota mediated communication with the bone marrow.

The results presented here suggest a model whereby gut colonization with *C. scindens* increases serum deoxycholate that then acts on the marrow to increase transcription of genes that support granulocyte monocyte progenitor (GMP) expansion, such as CCAAT/enhancer-binding proteins CEBPA and CEBPB. Then, when a novel challenge occurs at a mucosal site (in this case infection with *E. histolytica*), a more robust neutrophil response results.

Future studies will examine the precise mechanisms by which *C. scindens* colonization alters bone marrow hematopoiesis, which are not fully elucidated by these studies. However this work yields a mechanistic understanding of how changes in the gut microbiome can result in antigen nonspecific protection from *Entamoeba histolytica* infection. The impact of the work extends beyond infectious disease to fundamental mechanisms of gut to bone marrow communication by commensal bacteria and innate trained immunity. These studies may help in development of novel treatments that modulate the severity of immune and inflammatory diseases by altering bone marrow production of inflammatory cells.

## Acknowledgments

We thank Tuhinur Arju, and Mamun Kabir at icddr, b, Jeremy Gatesman, Homer Ransdell, Alice Kenney and Sanford Feldman at the University of Virginia Center for Comparative Medicine, Michael Solga, Claude Chew, and Joanne Lannigan at the Flow Cytometry Core facility, AhnThu Nguyen at the Biology Department Genomics Core, and Katia Sol-Church and Alyson Prorock at the Genome Analysis and Technology Core, Todd Fox at the UVA Metabolomics core, and Epigentek, NY, for technical support.

## Author’s contributions

SLB, JLL, JU, NO, KW, MS, MS, NG, BAT and BM performed the experiments. SLB, JLL, DTA, ST, DO, NG, JZM, ZP, BM and BAT analyzed the data. WAP, SLB, JS, RH, JZM, supervised the experiments and data analysis. SLB and WAP developed the theoretical framework. All authors discussed the results and contributed to the preparation of the manuscript.

## Data and materials availability

Data is available in the manuscript and supplemental figures. Full sequencing data will be deposited to the GEO repository under accession number GSE121503, the SRA under accession number PRJNA503904 and under SRA and linked via the dbGaP accession number phs001478.v1.p1.

All data is available upon request.

## Supplementary Materials for

**Table ST1.** Serum DCA independently predicts intestinal *E. histolytica* infection in a childhood cohort in Bangladesh and fecal calprotectin improves the predictability of *E. histolytica* infection. The association of serum DCA with intestinal *E. histolytica* infection was evaluated in logistic regression, adjusting for HAZ at enrollment, monthly family expenditure, and mother’s education. The regression was performed with and without consideration of fecal calprotectin. Although the effect of fecal calprotectin on *E. histolytica* infection was only marginally significant, the model with fecal calprotectin showed better c-statistic and thus was preferred. The c-statistic of 0.896 for the final model indicated near excellent predictive power for intestinal *E. histolytica* infection. N= 40 children per condition.

| Risk factor | OR | 95% CI |  | p value |
| --- | --- | --- | --- | --- |
|  |  | Lower | Upper |  |
| Serum DCA | 0.19<sup>†</sup> | 0.09 | 0.40 | <0.0001 |
| Fecal calprotectin | 0.95<sup>†</sup> | 0.90 | 1.01 | 0.0978 |
| HAZ at enrollment | 0.86 | 0.47 | 1.59 | 0.6301 |
| Monthly family expenditure | 1.00<sup>†</sup> | 0.99 | 1.01 | 0.4728 |
| Mother's education | 0.28 | 0.06 | 1.26 | 0.0970 |

OR: odds ratio; CI: confidence interval; HAZ: height-for-age z-score; DCA: deoxycholic acid;
<sup>†</sup> OR was calculated for 100 unit increment in the risk factor

**Figure S1.**
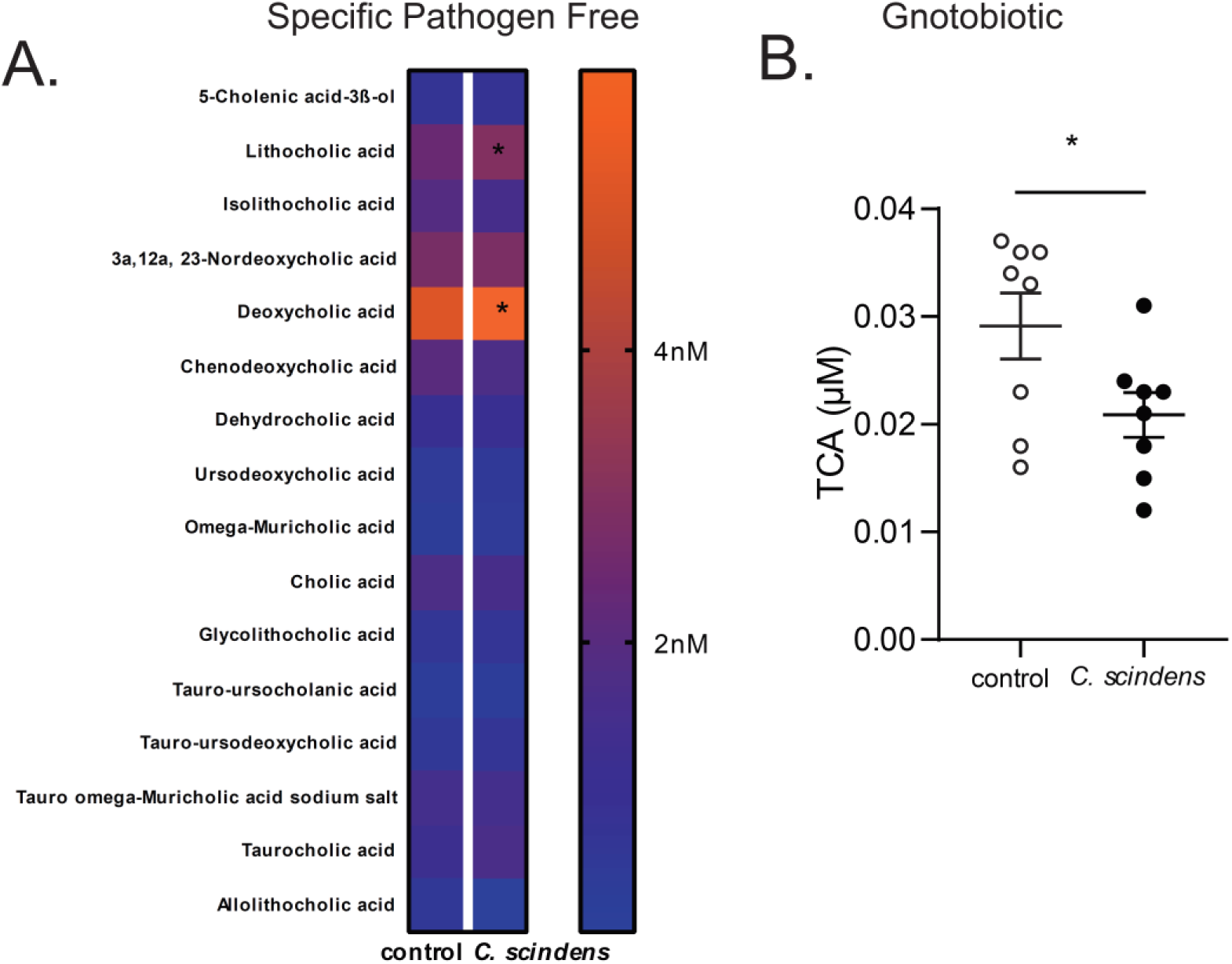
Serum bile acid profile of *Clostridium scindens* colonized mice is associated with increased bile acid deconjugation in both conventionally raised and gnotobiotic mice. CBA/J mice were colonized with *C. scindens* (ATCC® 35704), a bacterium capable of the 7α-dehydroxylation of bile acids, over three weeks prior to intracecal infection with *E. histolytica*. **(A)** Serum bile acids were measured using ultra-performance liquid chromatography-mass spectrometry in *C. scindens* gavaged and control animals. *= p<0.05, Fishers LSD. **(B)** C57BL/6 germ free mice were colonized with *C. scindens* with a single gavage. Serum bile acids were measured using ultra-performance liquid chromatography-mass spectrometry in *C. scindens* gavaged and control animals. *= p<0.05, Students T test, two tailed. Significant results with taurocholic acid (TCA) only. N =8 mice per group

**Figure S2.**
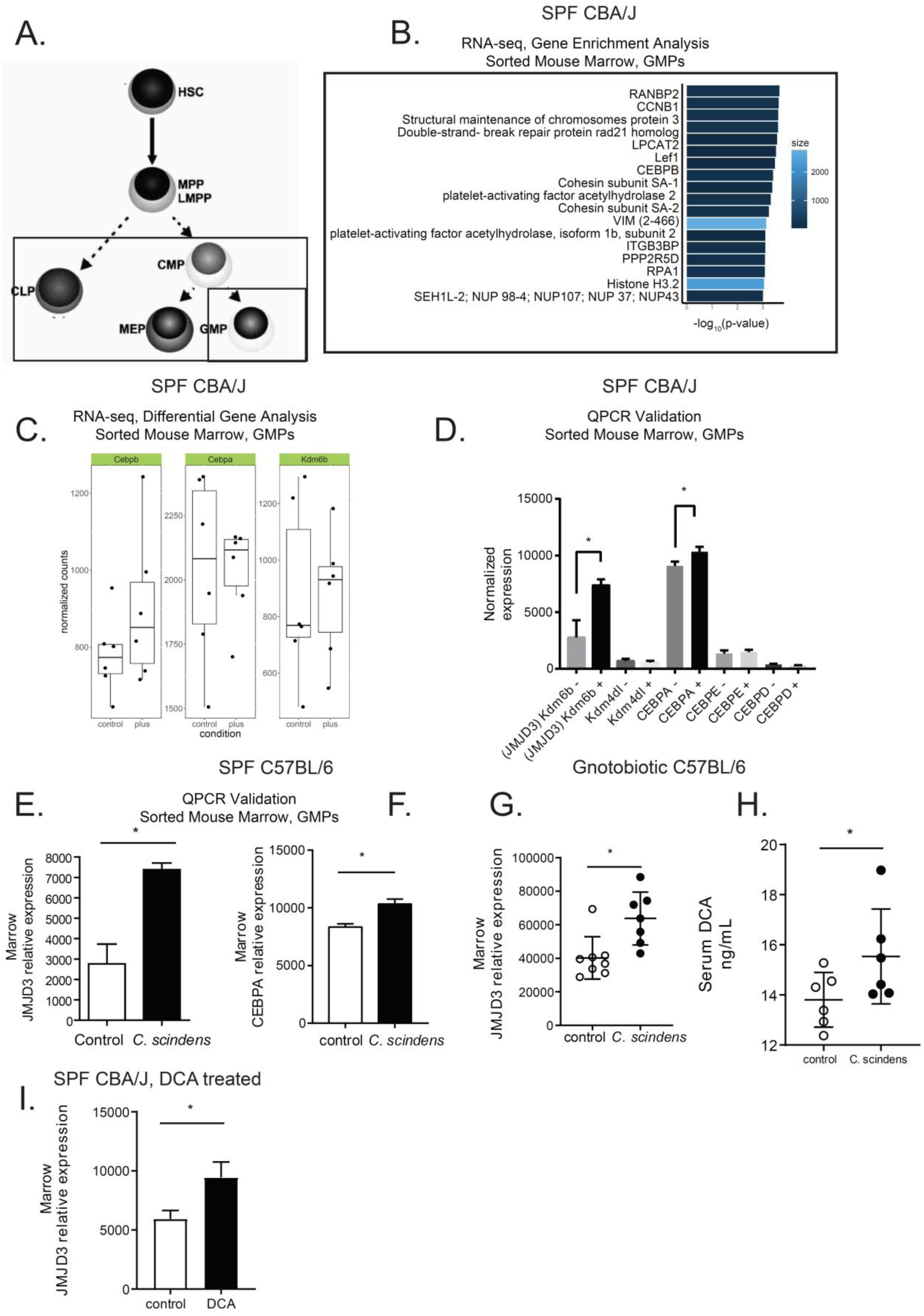
Intestinal colonization with *Clostridium scindens* increases expression of H3K27 demethylase JMJD3 and granulopoiesis promoting transcription factors in bone marrow granulocyte monocyte progenitors. (A) Bone marrow was sorted into CMP, GMP, MEP and CLP from specific pathogen free (SPF) CBA/J **(B, C, D, I)**, C57BL/6 **(E,F)** or C57BL/6 Gnotobiotic mice **(G,H)** that were colonized with *Clostridium scindens* or treated with DCA **(I). (B, C)** RNA-seq analysis and **(D-G, I)** qPCR was performed to validate enriched genes from RNA-seq on cDNA prepared from RNA isolated from sorted GMPs. **(B)** Gene set enrichment results obtained using ConsensusPathDB. The plot shows the most over-represented functional gene clusters associated with the top 50% of genes ranked by unadjusted P value. Note that the CEBPB gene set was identified as an enriched functional network using this unbiased approach. (**D-G, I)** Expression of noted genes was normalized to a housekeeping gene (S14). **(G)** Serum DCA was measured via ELISA **(H).** *= p<0.05, Student’s t-test, One Way ANOVA with Tukey posttest. N =6-12 mice per group

**Figure S3.**
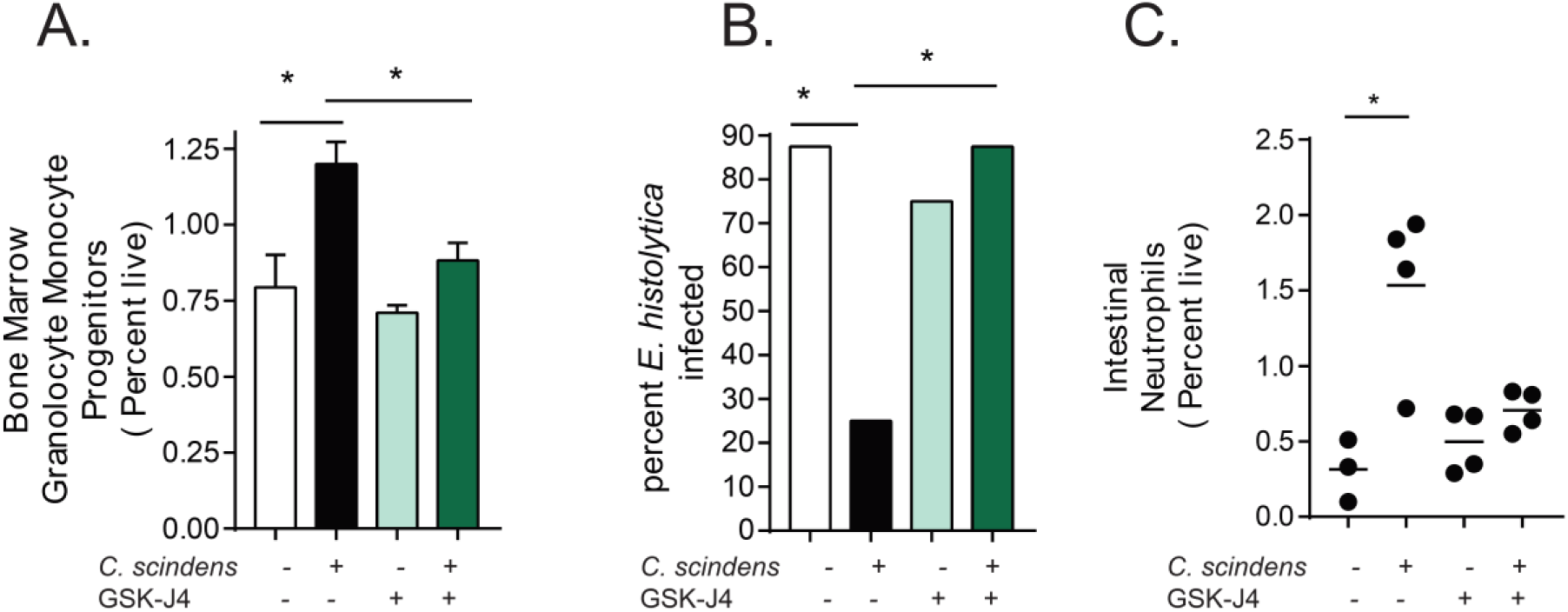
Blockade of H3K27 demethylase JMJD3 during *C. scindens* colonization abrogates marrow GMP expansion and intestinal protection from *E. histolytica.* CBA/J mice were treated with an inhibitor of the epigenetic mediator JMJD3 (GSK-J4) (**-**, **+**) before and during *C. scindens* colonization (**-**, **+**) but prior to infection with ameba. **(A)** Bone marrow progenitor populations, **(B)** percent infectivity and **(C)** intestinal neutrophils were analyzed via flow cytometry and culture. *= p<0.05, Student’s t-test, Mann–Whitney U test, one way ANOVA with Tukey posttest. N =4-12 mice per group.

**Figure S4.**
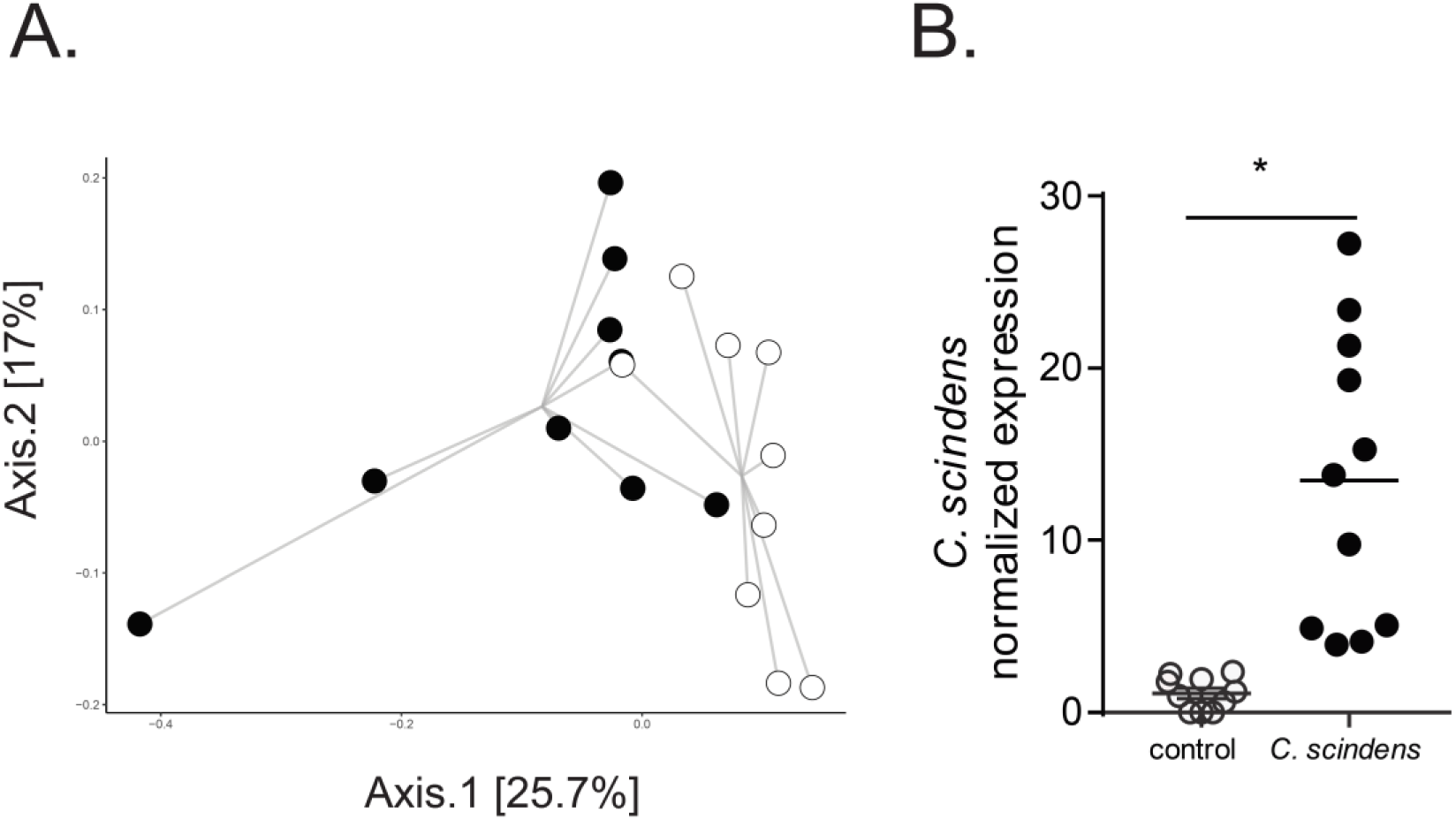
Introduction of *Clostridium scindens* to the gut microbiota alters community structure in mice. CBA/J mice were colonized with bile acid 7α-dehydroxylating bacteria *C. scindens* (ATCC® 35704) over three weeks prior to intracecal infection with *E. histolytica.* **(A)** Composition of the cecal microbiota community structure was determined by sequencing of V4 region of the 16S rRNA gene. PCoA of Bray-Curtis dissimilarities, the groups are significantly different by PERMANOVA, *p* = 0.002. **(B)** Relative expression of *C. scindens* baiCD stereo-specific 7alpha/7beta-hydroxy-3-oxo-delta4-cholenoic acid oxidoreductase normalized to total eubacterial 16S rRNA was determined via qPCR. *= p<0.05. N =9-12 mice per group.

**Figure S5.**
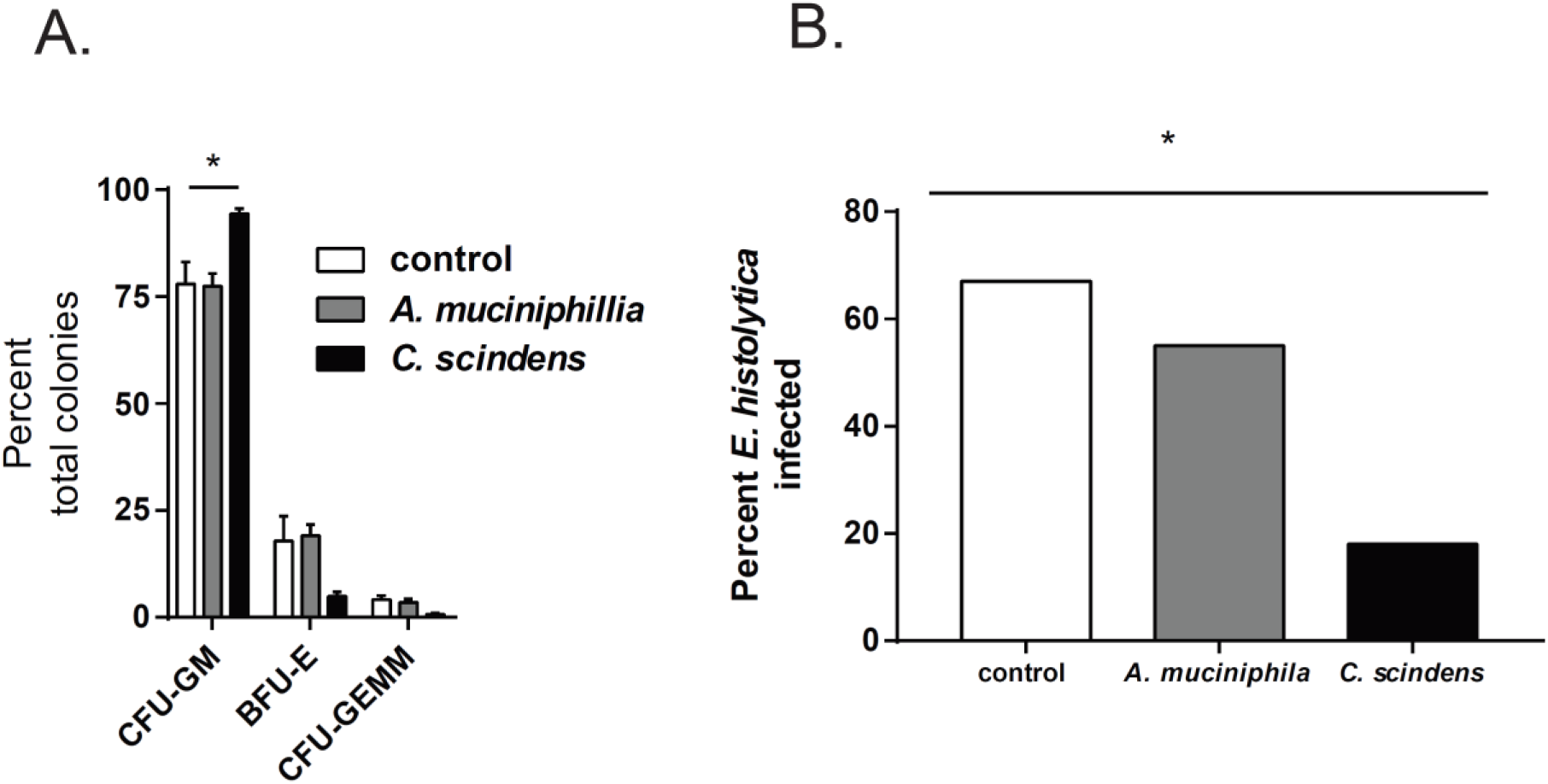
Introduction of human commensal *Akkermansia muciniphila* to the gut microbiota does not increase marrow GMPs or provide significant protection from *E. histolytica.* (A) CBA/J mice were colonized with *Akkermansia muciniphila* (ATCC ®BAA-835) or *C. scindens* (ATCC® 35704) over three weeks prior to intracecal infection with *E. histolytica*. **(B)** Composition of marrow hematopoietic precursors was determined via colony forming assays in mice. Percent infectivity with ameba at day six following infection was determined via cecal culture in trophozoite supporting media. *= p<0.05, Student’s t-test, bars and error bars are mean and SEM. N =6 mice per group.

## Methods

### Mice

Five week old male CBA/J mice (Jackson Laboratories or RAG 1 KO mice (Jackson Laboratories) were housed in a specific pathogen–free facility in micro isolator cages and provided autoclaved food (Lab diet 5010) and water ad libitum. Specific pathogen free status was monitored quarterly. Quarterly, a sentinel mouse (or rat) was removed from each room and humanely euthanized for serologic evaluation, examination of pelage for fur mites, and examination of cecal contents for pinworms. The serologic assays, conducted in-house using CRL reagents, were MHV, EDIM, GD-7, MVM, MPV, and MNV, (Sendai, PVM, RPV/ KRV/H-1, *M. pulmonis*, and SDAV for rats). In the final quarter, a comprehensive serology was run which included the above agents plus K-virus, MCMV, MTV, LCM, Ectromelia, Polyomavirus, Reovirus-3, and mouse adenoviruses (K87 and FL). For Gnotobiotic experiments Germ free C57BL/6 mice (Taconic) were housed in flexible film units (Park Bio) in a facility regularly monitored for germ free status by aerobic and anerobic culture and the above assays by the UVA Center for Comparative medicine. Cylinders were prepared and supplies and sterile food and water were introduced into the units according to SOP. Germ-free mice from Taconic were introduced into the units. Mice were gavaged once with sterile media or *C. scindens* as in SPF mice below. Aerobic and anerobic cultures of fecal samples were performed at introduction and once during the 2 week experiment as well as Sanger sequencing of isolated stool DNA using broad range eubacterial primers to confirm mono-association with *C. scindens* and germ free status of control animals. EUB forward 5’-ACTCCTACGGGAGGCAGCAGT-3’EUB reverse 5’-ATTACCGCGGCTGCTGGC-3’. One-week and two weeks post animal placement in the unit, surface cultures were performed to ensure sterility of unit. All contact areas, shipping containers and transfer apparatus were also cultured to confirm germ free status. All procedures were approved by the Institutional Animal Care and Use Committee of the University of Virginia. All experiments were performed according to provisions of the USA Animal Welfare Act of 1996 (Public Law 89.544). All experiments shown are representative of 2-4 experimental replicates.

### Clostridium scindens colonization

CBA/J mice (Jackson) were colonized with bile acid 7α-dehydroxylating bacteria *C. scindens* (ATCC® 35704) over three weeks prior to intracecal infection with *E. histolytica* or analysis for gut microbiome community structure, or marrow RNA seq and ChIP. Mice were gavaged with 100ul of overnight culture at an optical density of 1.4 at 600nm or media control (BHI, Anerobe Systems, AS-872) once per week for specific pathogen free mice over three weeks, and once for C57BL/6 (Taconic) gnotobiotic mice over two weeks.

### Intravenous deoxycholate treatment

CBA/J mice were treated intravenously via tail vein injection by a trained veterinary technician 3 times a week, over two weeks with 400uL of PBS or 400uL of 0.20mg/mL DCA (Sigma) in PBS per treatment. Mice were healthy during DCA administration and no liver damage as measured via serum ALT ELISA (Cloud-Clone Corp, SEA207Mu) was observed. Liver also appeared grossly normal on dissection.

### Adoptive Marrow Transplant

Donor and recipient CBA/J mice were colonized with bile acid 7α-dehydroxylating bacteria *C. scindens* (ATCC® 35704) or treated with media controls as above. CBA/J mice colonized with *C. scindens* (+) or not were lethally irradiated, then immediately given whole marrow from *C. scindens* (+) or *C. scindens* (-) donors then allowed to recover for 7 weeks prior to ameba challenge. Irradiation was performed with 900 Rad in a single dose from a Shepard irradiator, Mark 1 Model 68A Dual, Serial Number 1163 with a Cs-137 source. All irradiation experiments were supervised by a member of the University of Virginia Environmental Health and Safety team. Mice were place on placed on sulfamethoxazole-trimethoprim containing water 3 days prior and 21 days post bone marrow transplant. Experiments shown are representative of 2 experimental replicates.

### *E. histolytica* culture and intracecal injection

Animal-passaged HM1:IMSS *E. histolytica* trophozoites were cultured from cecal contents of infected mice in complete trypsin-yeast-iron (TYI-33) medium supplemented with Diamond Vitamin mixture (JRH Biosciences), 100 U/ml of both penicillin and streptomycin, and 5% heat inactivated bovine serum (Sigma-Aldrich). Prior to injection, trophozoites were grown to log phase, and 3 x10^6^ parasites were suspended in 100 µL culture media and injected intracecally (6). Data analysis and graphing was performed with Graphpad Prism 8.0. Final figures were modified and arranged in Adobe Illustrator.

### JMJD3 Blockade

Mice were treated intraperitoneally with hybridoma grade DMSO in PBS (Sigma) or GSK-J4 in DMSO/PBS (100uL, DMSO/PBS, 25 mg/kg, Cayman Chemical #12074) one day before and during *C. scindens* colonization (once per week) but not during *E. histolytica* infection. Experiments shown are representative of 2 experimental replicates.

### Flow cytometry of intestinal cells

Minced intestinal tissue was digested in Liberase TL (0.17 mg/ml Roche) and DNase (0.5 mg/ml, Sigma) for 45 min at 37°C and processed into a single cell suspension following washing with a buffer containing EDTA. 1×10^6^ cells per mouse were stained with antibodies from Bio Legend, CD11c-BV421, CD4-BV605, Ly6c-FitC, CD3e-PerCp CY 5.5, SIGLEC F-PE, Ly6G PE Cy7, CD11b-APC, CD8a AF700, CD45-APC Cy7. Flow cytometric analysis was performed on an LSR Fortessa (BD Biosciences) and data analyzed via FlowJo (Tree Star Inc.). All gates were set based on fluorescence minus one (FMO) controls. Further data analysis and graphing was performed with Graphpad Prism 7.0. Final figures were modified and arranged in Adobe Illustrator. Experiments shown are representative of 2-4 experimental replicates.

### Bone marrow flow cytometry and cell sorting

Bone marrow cells were isolated from femur, fibia and tibia of mice by centrifugation in custom made microcentrifuge tubes composed of a 0.5ml microcentrifuge tube with a hole punched in the bottom nested inside a 1.5mL microscentrifuge tube (VWR). Bone marrow cells were stained with PerCP-Cy5.5-labeled lineage (Lin) markers (TCRb, CD3e,CD49b, B220Gr1, CD11c, CD11b), anti-CD34-Brilliant Violet 421, c-Kit-Brilliant Violet 605, CD127-PE-Cy7, FcgRII-III (CD16/CD42)-APC-Cy7, and Sca-1-APC, CD150 PE, Live dead staining with Zombie Aqua (Bioledgend) or 7AAD for sort experiments. Flow cytometric analysis was performed on an LSR Fortessa (BD Biosciences) or cells sorted on a Becton Dickinson Influx Cell Sorter into RNA later (Qiagen) or Cryostore CS10 (Stemcell) with 7AAD as a live dead stain. Data analyzed via FlowJo (Tree Star Inc.). All gates were set based on fluorescence minus one (FMO) controls. Common Lymphoid Progenitors (CLP) are Lin-IL-7R+c-Kit^int^Sca-1^int^, Common Myeloid Progenitors (CMP) are Lin-c-Kit+Sca-1-CD34+FcgRII-III^int^; Granulocyte-Monocyte-Progenitors (GMP) are Lin-c-Kit+Sca-1-CD34+FcgRII-III^hi^; Megakaryocyte–Erythroid Progenitors (MEP) are Lin-c-Kit+Sca-1-CD34-FcgRII-III-. Experiments shown are representative of 2-4 experimental replicates.

### ChIP-qPCR

For ChIP-QPCR experiments approximately 6,000 GMPs from *C. scindens* colonized or control mice (CBA/J) were sorted on an Influx cell sorter into Cryostore CS10 (Stemcell) as above. The samples were thawed then subjected to chromatin isolation, chromatin shearing, DNA isolation, and ChIP. ChIP-qPCR and data analysis was then preformed. *Chromatin Isolation:* Chromatin was isolated using the ChromaFlash Chromatin Extraction Kit (EpiGentek, Cat. #P-2001). *Chromatin Fragmentation:* Chromatin was sheared using the EpiSonic 2000 Sonication System (EpiGentek, Cat. #EQC-2000) based on the standard operation protocol for small amounts of cells. Chromatin was sonicated for 20 cycles with 45” On and 15” Off. *Chromatin Quantification:* Total volume of chromatin solution was 60 μl for each sample. The sheared chromatin concentration was measured by fluorescence quantification of chromatin associated DNA. *DNA Purification:* DNA was purified using 5 μl of sheared chromatin as input. DNA was eluted with 15 μl of water. 1 μl of purified DNA was used for fluorescence quantification. *Antibody validation:* The anti-H3K4me3 (Epigentek Cat. #A-4033), anti-H3K27me3 (Epigentek Cat. #A-4039) antibodies were validated using the Pre-Sure ChIP Antibody Validation Kit (EpiGentek, Cat. #P-2031). The ChIP-Grade Intensity (CGI) is 4.0 for H3K4me3, and 4.2 for H3K27me3, respectively. *Chromatin Immunoprecipitation:* Because of small amount of chromatin for each sample, the ChIP reaction was based on the high sensitivity ChIP protocol modified from P-2027 kit. 50 ng of sheared chromatin samples in 200 μl ChIP assay buffer were added into the wells coated with 0.5 μg of anti-H3K4me3 or anti-H3K27me3, respectively. 2 ug of Jurkat cell chromatin was used as a positive control. The samples were incubated at room temperature for 180 minutes with continuous shaking (100 rpm). After incubation, the wells were washed and the chromatin immunoprecipitated DNA was purified and eluted in 14 μl of water. *qPCR:* was performed in duplicate using 1μl of DNA and gene-specific primers designed for the target gene region for 60 cycles. Un-ChIPed DNA (10%) was used as input for determining enrichment efficiency (Input%). Primer sequences based on the targeted CEBPA and CEBPB region sequences are listed below/ CEBPA-1: CEBPA-TSS; CEBPA-2: CEBPA-promoter; CEBPB-1: CEBPB-TSS; CEBPB-2: CEBPB-promoter

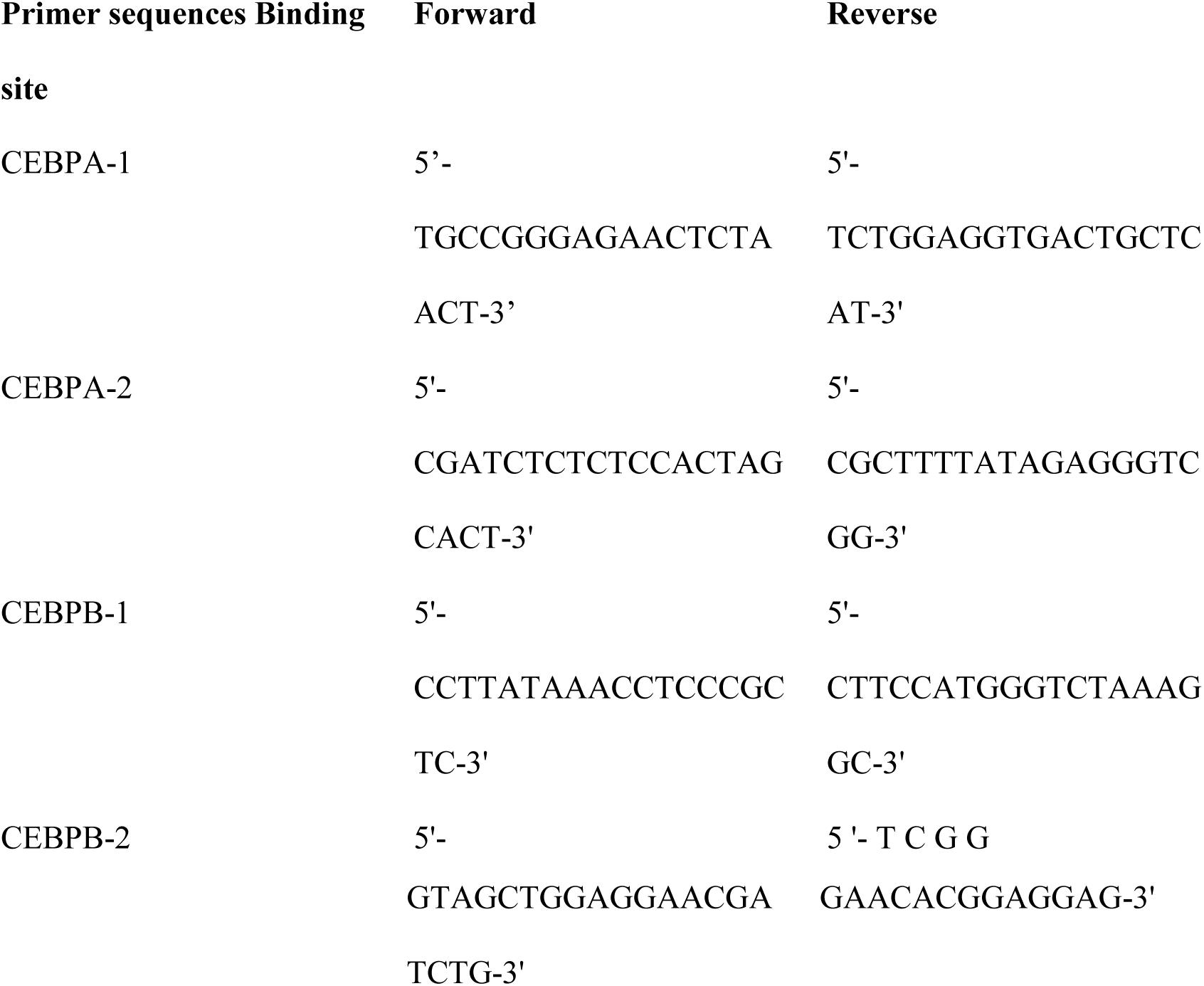

### Colony forming assay for determination of bone marrow hematopoietic precursors

Bone marrow cells were isolated (*33*) and then cultured in methylcellulose-based medium that included, 3 units/mL Epo, 10 ng/mL mouse recombinant IL-3, 10 ng/mL human recombinant IL-6, and 50 ng/mL mouse recombinant stem-cell factor per manufacturer procedures (M3434; StemCell Technologies, Vancouver, BC). Colony formation of burst-forming unit–erythroid (BFU-Es), colony-forming unit–granulocyte/monocyte (CFU-GMs), and CFU granulocyte/erythrocyte/monocyte/macrophage (CFU-GEMMs) were analyzed after 7 days. Experiments shown are representative of 2 experimental replicates.

### RNA sequencing and data analysis

GMPs were isolated from six control and six *C. scindens-*treated mice as described above. RNA was isolated from approximately 7,000 sorted GMPs per mouse utilizing the Qiagen RNeasy Micro Kit. Ribosomal RNA depletion was performed using the NEBNext® rRNA Depletion Kit (Human/Mouse/Rat) and alternate protocol for low yield RNA. Directional cDNA libraries were generated the NEBNext® Ultra™ Directional RNA Library Prep Kit for Illumina®, including 15 PCR cycles of amplification. Multiplexed samples were sequenced (75 bp paired-end reads) using the NextSeq 500 platform. For sequencing, libraries were sequenced with the NGS NextSeq kit - 150 cycle High Output Kit, paired end 75×75 bp read. Data analysis was performed first by the UVA Bioinformatics Core. Reads were mapped to the mouse transcriptome (GRCm38) using Salmon and gene level abundances quantified using tximport. Differential gene expression analysis was then performed in R using DESeq2(*34*), which yielded no differentially expressed genes with FDR-corrected p-values < 0.05. In a second round of analysis, RNA-seq data were analyzed by aligning the raw reads to the mouse mm10 version of the genome using HISAT2(*35*) (v2.0.4) and transcripts were assembled and quantified using StringTie (v1.3.4d)(*36*) and GENCODE vM19 annotations. Using this second approach, a small collection of 20-30 significantly affected genes were identified. ConsensusPathDB (*37*) was used to identify functional enrichment in the top 50% of genes ranked by unadjusted P value. RNA-seq data are available from the Gene Expression Omnibus (accession number GSE121503).

### 16S rRNA Gene Amplicon Sequencing (Mouse)

DNA was isolated from mouse cecal lysate (QIAamp DNA Stool Mini Kit). The V4 region of the 16S rRNA gene was amplified from each sample using the dual indexing sequencing strategy as described previously(*38*). Sequencing performed on the Illumina MiSeq platform, using a MiSeq Reagent Kit V2 500 cycles (Illumina cat# MS102-2003), according to the manufacturer’s instructions with modifications found in the Schloss SOP: https://github.com/SchlossLab/MiSeq_WetLab_SOP. The mock community produced ZymoBIOMICS Microbial Community DNA Standard (Zymo Research cat# D6306) was sequenced to monitor sequencing error. The overall error rate was 0.02% as determined using the software package mothur version 1.39.5 following the Illumina MiSeq standard operating procedure (*39*).

### 16S rRNA Gene V4 region sequencing, sample selection and extraction (Human)

Diarrheal and non-diarrheal reference stools were collected during scheduled study visits (scheduled visits took place at enrollment and at 6, 10, 12, 14, 17, 18, 39, 40, 52, 53, 65, 78, 91, 104 weeks of age)(*19*). Samples were brought into the study clinic and stool was transported from the field to our laboratory at 4°C, aliquoted in DNase- and RNase-free cryovials, and stored at −80°C on the day of collection. 200ug was removed for total nucleic acid extraction. Positive extraction controls were achieved by spiking phocine herpesvirus (Erasmus MC, Department of Virology, Rotterdam, The Netherlands) and bacteriophage MS2 (ATCC 15597B; American Type Culture Collection, Manassas, VA) into each sample during the extraction process. The fecal DNA was then tested for *E. histolytica* by use of a multiplex qPCR assay to detect parasitic protozoans as described by Haque et al (38). DNA Samples positive for *E. histolytica* and non-diarrheal reference samples were shipped to our laboratory at UVA for library construction.

### Library construction for next-generation sequencing (Human)

The entire 255bp V4 region of the 16 S rDNA gene was amplified as previously described(*40*), using phased Illumina-eubacteria primers to amplify the V4 16 S rDNA region (515F – 806R) and to add the adaptors necessary for illumina sequencing and the GOLAY index necessary for de-multiplexing after parallel sequencing. Negative controls included the addition of extraction blanks that were tested throughout the amplification and sequencing process to ensure they remained negative. As a positive PCR control, DNA extracted from the HM-782D Mock Bacteria Community (ATCC through BEI Resources) was run on each plate and added to the library. The library was then sent to UVA Biomolecular core facility. A PhiX DNA library was spiked into the 16S sequencing run (20%) to increase genetic diversity prior to parallel sequencing in both forward and reverse directions using the Miseq V3 kit and machine (per manufacturer’s protocol).

### 16S rRNA Gene Amplicon Curation and Analysis

All 16S data curation and analysis was performed using R version 3.5.1. Sequences were curated using the R package DADA2 version 1.10.1, following the DADA2 pipeline tutorial v1.8 (*41*). Briefly, reads were filtered and trimmed using standard parameters outlined in the DADA2v1.8 pipeline. The error rates for the murine or human amplicon datasets were determined using the DADA2’s implementation of a parametric error model. Samples were then dereplicated and sequence and variants were inferred. For the 16S data from murine samples, overlapping forward and reverse reads were merged and sequences that were shorter than 250bp or longer than 254bp were removed. For the 16S data from human samples, only forward reads were used. Finally, chimeras were removed. Taxonomy was assigned to amplicon sequence variants (ASVs) using the DADA2-formatted SILVA taxonomic training data release 132 (*42*). A partial sequence from *[Lachnoclostridium] scindens ATCC 35704* (NCBI Reference Sequence: NR_028785.1) was added to the Silva training data v132 to attempt to identify *Clostridium scindens* ASVs. Following sequence curation, the packages phyloseq v1.26.1(*43*), vegan, dplyer and ggplot2 were used for analysis and generation of figures. This includes determining the axes for the PCoA plots of Bray-Curtis dissimilarities (beta-diversity) calculated from rarified sequence abundance. Additionally, the package vegan was used to determine significant differences between groups with PERMANOVA. The sequences associated with analysis of the murine data were deposited to the SRA under the PRJNA503904. The sequences associated with analysis of the human data will be deposited to the SRA and linked via the dbGaP accession number phs001478.v1.p1. Full details of the design of the human cohort study have been described (*19*) and all studies were approved by the Ethical Review Committee of the ICDDR, B and the Institutional Review Boards of the Universities of Virginia and informed consent was obtained after the nature and possible consequences of the studies were explained in all cohort studies. Final figures were modified and arranged in Adobe Illustrator CC.

### *C. scindens* culture, marrow and cecal lysate qPCR

Purity of *C. scindens* culture was confirmed via qPCR and Sanger sequencing with broad specificity eubacteria primers, EUB forward 5’-ACTCCTACGGGAGGCAGCAGT-3’EUB reverse 5’-ATTACCGCGGCTGCTGGC-3’, and *C. scindens* baiCD stereo-specific 7alpha/7beta-hydroxy-3-oxo-delta4-cholenoic acid oxidoreductase primers, BaiCD F - 5′-CAGCCCRCAGATGTTCTTTG-3′ BaiCD R - 5′-GCATGGAATTCHACTGCRTC-3′ *C. scindens* colonization was measured via qPCR from cecal lysate (QIAamp DNA Stool Mini Kit). qPCR for baiCD with SYBR green was performed and data were normalized to expression of a conserved *Eubacteria* 16s RNA gene (EUB) (*44*). Primer concentrations, annealing temperatures, and cycle number were optimized for each primer pair. For each primer pair, a dilution curve of a positive cDNA sample was included to enable calculation of the efficiency of the amplification. The relative message levels of each target gene were then normalized to EUB or the mouse housekeeping gene S14 using a method described and utilized previously (*12, 13, 45*). Data is presented as relative expression. For sorted marrow qPCR, RNA was isolated from approximately 7,000 sorted GMPs utilizing the Qiagen RNeasy Micro Kit. S14 forward 5’-TGGTGTCTGCCACATCTTTGCATC-3’, S14 reverse 5’ AGTCACTCGGCAGATGGTTTCCTT-3’,Jmjd3 forward 5’-CTCTGGAACTTTCATGCCGG-3’ Jmjd3 reverse, 5’-CTTAGCCCCATAGTTCCGTTTG-3,’ CebpA forward 5’-CAAAGCCAAGAAGTCGGTGGACAA, CebpA reverse 5’ – TCATTGTGACTGGTCAACTCCAGC CebpE forward 5’-TGTGGGCACCAGACCCTAAG, CebpE reverse 5’-GCTGCCATTGTCCACGATCT, CebpD forward 5’-CTTTTAGGTGGTTGCCGAAG, CebpD reverse 5’ GCAACGAGGAATCAAGTTTCA, Kdm4dl forward 5’-CATGGTCACCTTTCCCTATGG, Kdm4dl reverse 5’-AAAATTGATGGCCTCTGCG. Primers were purchased from Integrated DNA Technologies Coralville, Iowa, USA.

### Serum deoxycholate ELISA

Serum Deoxycholate was measured via ELISA (Cloud-Clone Corp. CES089Ge) in 80 children each from two birth cohorts in Mirpur Dhaka, Bangladesh. Full details of the design of these two birth cohort studies, including socioeconomic status data, have been described (*19, 46*) and all studies were approved by the Ethical Review Committee of the ICDDR, B and the Institutional Review Boards of the Universities of Virginia and informed consent was obtained after the nature and possible consequences of the studies were explained in all cohort studies. Children were approximately two years of age and serum was selected by identifying diarrheal stools within 6 months of the blood draw that were *E. histolytica* positive (n=40) and negative (n=40) as identified via qPCR in both cohorts (*47*). Association of serum DCA with intestinal *E. histolytica* infection was evaluated in a logistic regression, adjusting for HAZ at enrollment, monthly family expenditure, and mother’s education. The regression was performed with and without consideration of fecal calprotectin. Improvement of model fit with addition of calprotectin was evaluated by log-likelihood ratio test. The model with fecal calprotectin showed a slightly improved model fit, in which c-statistic of 0.896 indicated near excellent predictive power for intestinal EH infection. Serum was measured in 6 mice from each of at least two experiments that were gavaged with *C. scindens* or media control as described utilizing the same kit or in DCA treated mice. For both children and mice 25uL of serum was utilized at a 1:2 (humans) or 1:4 (mice) dilution following kit protocol with a 10 minute development step following administration of substrate solution. The logistic regression analyses were performed using the function “glm” in R software version 3.5.2 (r-project.org), while model fitting and predictability were evaluated using the R packages “lmtest” and “pROC” respectively.

### Targeted profiling of serum bile acids using UPLC-MS

In specific pathogen free mice, Plasma bile acids were quantified as previously described (*48*) by ACQUITY ultraperformance liquid chromatography (UPLC) (Waters, Ltd., Elstree, UK). Briefly, 100 μl of plasma were spiked with isotopically labeled bile acid standards followed by the addition of 300 μl of ice-cold methanol to facilitate protein precipitation. Bile acids were separated over a 15-minute gradient on an ACQUITY BEH C8 column (1.7 μm, 100 mm x 2.1 mm), detected by a Xevo TQ-S mass spectrometer (Waters, Manchester, UK) operating in the negative ionization mode (ESI-) and assayed using multiple reaction monitoring (MRM). In gnotobioic mice Plasma bile acids were measured using the Biocrates Bile Acids kit (Biocrates Life Sciences AG, Innsbruck, Austria), which detects 20 bile acids. Briefly, 10ml of murine plasma samples was added to filter plates and samples were processed according to manufacturer’s protocol. All reagents used during sample preparation were UHPLC-MS grade. Samples were analyzed by LC-MS/MS on a Waters (Milford, MA) I-Class Acquity chromatography system in-line with a Waters TQ-S mass spectrometer. Compounds were analyzed using TargetLynx XS software with the results subsequently imported into MetIDQ software (Biocrates) for quality control (QC) validation. Raw metabolite concentrations (mM) were normalized using identical QC samples across the plate to control for variation over the course of the run. QC samples were proprietary samples spiked with metabolites measured during targeted metabolomics. Normalized metabolite concentrations were exported for further analysis and screened for inclusion in the data set based on two criteria. First, metabolites must be within the valid range in at least 66% of quality control samples run on a plate to be included in analysis. Second, metabolites must be above the limit of detection in at least 60% of all measurements to be included in the valid data set. Data set validation was performed using R software version 3.4.3 (r-project.org)

